# Comprehensive valuation of the ecosystem services of the Arctic National Wildlife Refuge

**DOI:** 10.1101/2020.03.09.983999

**Authors:** Adam C. Turner, Margaret A. Young, Maureen R. McClung, Matthew D. Moran

## Abstract

Ecosystem services (ES) have been well studied in most biomes, but the Arctic tundra has received little attention, despite covering over 10% of terrestrial Earth. Using established ES methodologies, we calculated values for the United States Arctic National Wildlife Refuge, a region virtually undisturbed by humans, but slated for future oil and gas drilling. We estimated the Refuge is worth about 1,709 USD/hectare/year, equal to over 13 billion USD annually.

Globally important services, such as climate regulation (e.g., carbon storage) and non-use services (e.g., aesthetic information), contributed the most value and were similar to valuations from more productive ecosystems. Local services made smaller contributions to the total, but they remain vitally important to local indigenous cultures. Strikingly, a contingent valuation survey of U.S. residents found that, after neutral educational information, willingness-to-pay to maintain the Refuge in its current state exceeded estimated values of the oil and gas deposits.

Our study shows that citizens may value Arctic habitats beyond their traditional economic development potential. Our comprehensive ecosystem services valuation suggests that maintaining the Refuge in its current condition (i.e., *de facto* wilderness) with its full range of ES is more valuable to humanity compared to development for oil and gas.

## Introduction

Ecosystem services (ES) assessments place an economic value on resources in natural landscapes that are beneficial to humans. In recent decades, most biomes have had multiple ES values estimated (Costanza et al. 1997; Costanza et al. 2014; de Groot et al. 2012) with global annual ecosystem services values calculated at over $100 trillion dollars a year (de Groot et al. 2012; Costanza et al. 2014). However, two biomes that cover a large proportion of the terrestrial surface remain relatively understudied: desert and tundra (Malinauskaite et al. 2019). Taylor et al. (2017) developed a valuation for a portion of the Chihuahuan Desert and O’Garra (2017) has provided an estimate of some key ecosystem services for Arctic tundra. Considering that Arctic tundra alone covers 5.7 million km^2^ (Shaver et al. 1992) of the earth’s surface and, by one estimate, contains 50% of the global belowground organic carbon (Tarnocai et al. 2009), it is likely that this biome contributes a large amount of ecosystem services (O’Garra 2017).

Arctic tundra is composed of relatively low productivity ecosystems, but these may still provide numerous ecosystem services on both a local and global scale. Tundra contains large stores of carbon, mostly located in the permafrost layer (Tarnocai et al. 2009), and is therefore vitally important for global climate regulation (Zimov et al. 2006). Productive fisheries occur in Arctic coastal territories (CAFF 2015), providing food both locally and for export (Zeller et al. 2011; Christiansen et al. 2014). Arctic tundra is largely undeveloped compared to other biomes (Watson et al. 2016), has significant wildlife populations (Johnson et al. 2005), and supports unique cultures (Hovelsrud et al. 2011), further increasing its likely contribution to a variety of information functions (e.g., recreational, aesthetic, cultural, scientific understanding, de Groot et al. 2002).

Global climate change and fossil fuel extraction threaten the arctic tundra biome across continents. As global temperature rises, the polar regions are increasing in temperature more rapidly than equatorial zones (Hansen et al. 2006, 2010), causing rapid environmental and social changes to these high latitudes (Wookey et al. 2009; Bhatt et al. 2013; Chapin et al. 2015).

Furthermore, oil and natural gas extraction is expanding into new areas of the Arctic, causing increased fragmentation, decline of ecosystem function, and negative impacts on wildlife (Boulanger et al. 2012; Johnson and Russell 2014) in areas that were previously some of the most intact ecosystems on Earth (Sanderson et al. 2002; Watson et al. 2016). As the globe continues to warm, Arctic areas are becoming more accessible to industrial development, especially by petroleum industries (Harsem et al. 2011). Considering that the circumpolar region is estimated to contain large quantities of oil and gas, up to 22% of the global reserves (Bird et al. 2008), enhanced fossil fuel extraction could further exacerbate climate change. Currently, there is little information on how ES relate to management of specific Arctic regions (Malinauskaite et al. 2019)

In the United States, Arctic tundra covers around one-third of the state of Alaska and plays an important role in the state’s economy. About 17% of the Alaskan tundra region is protected by the Arctic National Wildlife Refuge (Ibisch et al. 2016), which is one of the largest contiguous natural areas in the U.S. There is no road access to the Refuge, resulting in low human visitation. As such, the remoteness of the area provides a prime opportunity to study an undisturbed tundra landscape. The Refuge is also important to the subsistence, culture, and spiritual life of native people who live in and around the region. However, this area is under increasing threat from energy industries. Large oil and natural gas reserves are thought to occur in the Refuge (estimated at more than 10 billion barrels of oil; Attanasi 2005), mostly in the coastal plain region (known as the 1002 Area). When the Refuge was created in 1980, the U.S. Congress directed that additional natural and cultural resource studies occur in the 1002 area so that a future Congress could be better informed about potential oil and gas development. An intense public debate ensued with numerous unsuccessful attempts to open the area for drilling. More recently, the Tax Cuts and Jobs Act of 2017 (H.R.1 2018) was passed by the U.S. Congress and signed by the President, allowing for oil and natural gas exploration to proceed. By early 2020, no activity has yet commenced while environmental impact assessments are conducted.

Our primary goal in this study was to determine the ES values for the Arctic National Wildlife Refuge. Assuming the Arctic National Wildlife Refuge is representative of typical Arctic tundra habitat, this value can be added to the global ES estimate. This estimation could also be utilized to understand the impact of planned energy development. Since no ES estimates have been calculated for the Arctic National Wildlife Refuge, the true costs and benefits of planned oil exploration and production cannot be accurately determined. This ES value should be calculated prior to resource extraction so that we can determine if the ES value of the Arctic National Wildlife Refuge is greater or less than the economic benefits of fossil fuel development.

## Methods

### Study Site

The Arctic National Wildlife Refuge is located in the northeastern corner of Alaska and covers about 78,000 km^2^ (Fig. 1). It was established as a protected area in 1960 and expanded in 1980 under the Alaska National Interest Lands Conservation Act (Public Law 97-394 1980).

**Fig. 1.**
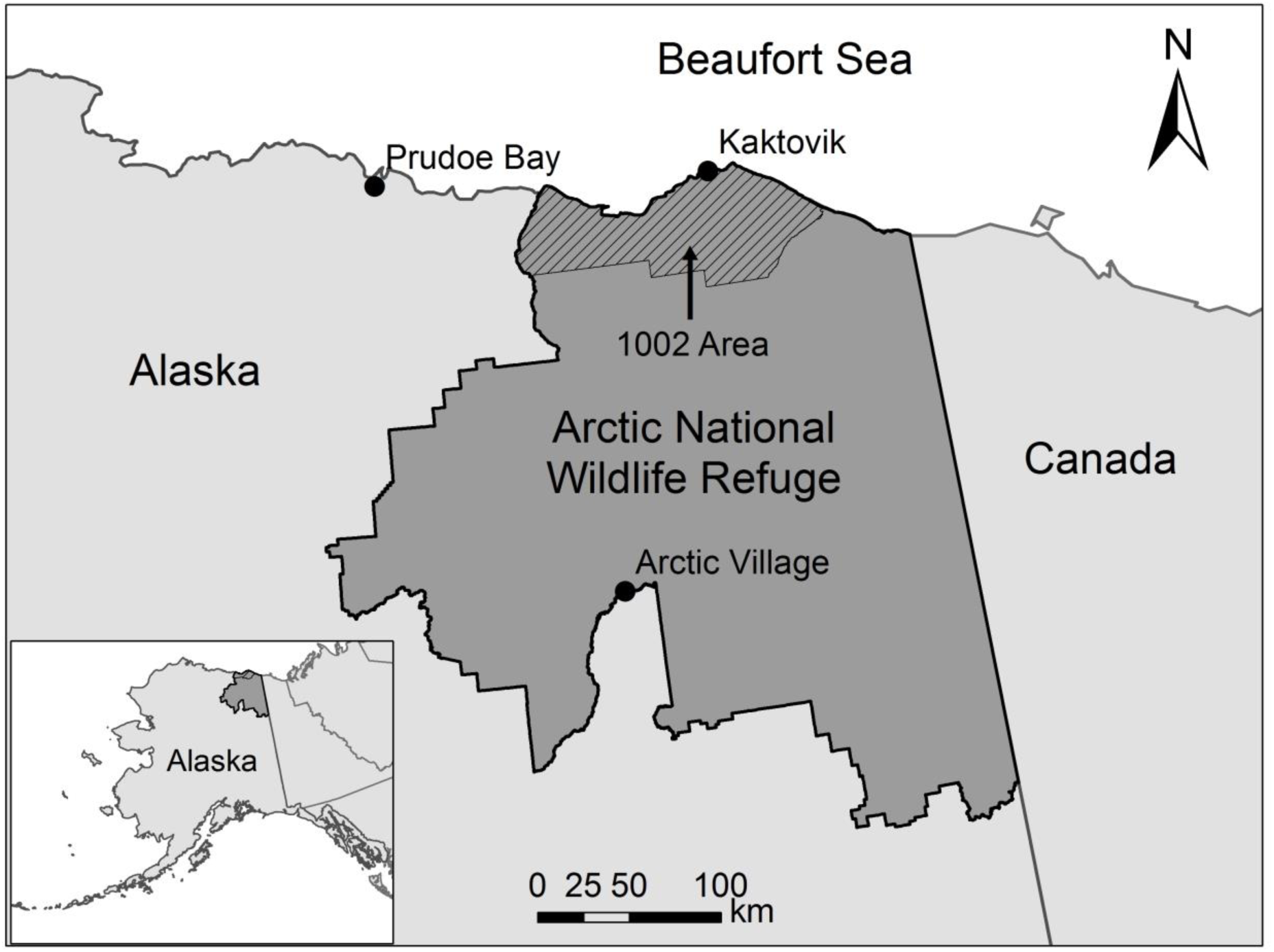
Map of the Arctic National Wildlife Refuge in relation to the 1002 area, Canada, and the U.S. state of Alaska (inset).

That law designated the 1.2 million acres of the coastal plain section (1002 Area) for potential oil and gas development, but only with future U.S. Congressional approval.

The Arctic National Wildlife Refuge is noted for its high Arctic biodiversity, roadless and wilderness characteristics, and large populations of animals (United States Bureau of Land Management 1986). It is particularly important for the Porcupine Caribou Herd (*Rangifer tarandus granti*), which calves annually on the coastal plain (Whitten et al. 1992), denning polar bears, and the large number of migratory birds which utilize the region for breeding (Douglas et al. 2002; Brown et al. 2007). Much of the Refuge is covered in typical Arctic tundra habitat, with smaller areas of boreal forest in the southern portions. It is considered by many scientists as one of the finest remaining Arctic habitats in the world (Brown 2002; National Research Council 2003).

Very few people live in the vicinity of the Refuge. The village of Kaktovik (population: 241) is located within the Refuge boundary along the Arctic coast and Arctic Village (population: 152) is located on the southern boundary of the Refuge (Fig. 1). Additionally, villages to the east in Canada, south along the Yukon River watershed, and southwest of the Refuge also utilize refuge resources. While these Alaska Native villages do not contain large population numbers, they depend very heavily on the subsistence and cultural resources of the Refuge.

### Estimating Ecosystem Services

We utilized the TEEB (2010) classification system, originally described in de Groot et al. (2002) as the model for classifying ecosystem services provided by biological systems (Fig. 2). While there are several different ES classification systems commonly utilized, there are broad similarities between each, making them roughly comparable (Costanza et al. 2017). We estimated each category of ES values using methods described in the literature (Costanza and Folke 1997; de Groot et al. 2012; Costanza et al. 2014, and described in detail below). We performed a comprehensive analysis of all the services established by de Groot et al. (2002) and TEEB (2010), recognizing that not all services are likely to be prominent in an Arctic environment. Since we had no *a priori* knowledge about which individual services were present, we attempted to calculate and report on all ES values regardless of their amount or explained why they were not found in this environment. All monetary estimates were inflation-adjusted to 2016 USD.

**Fig. 2.**
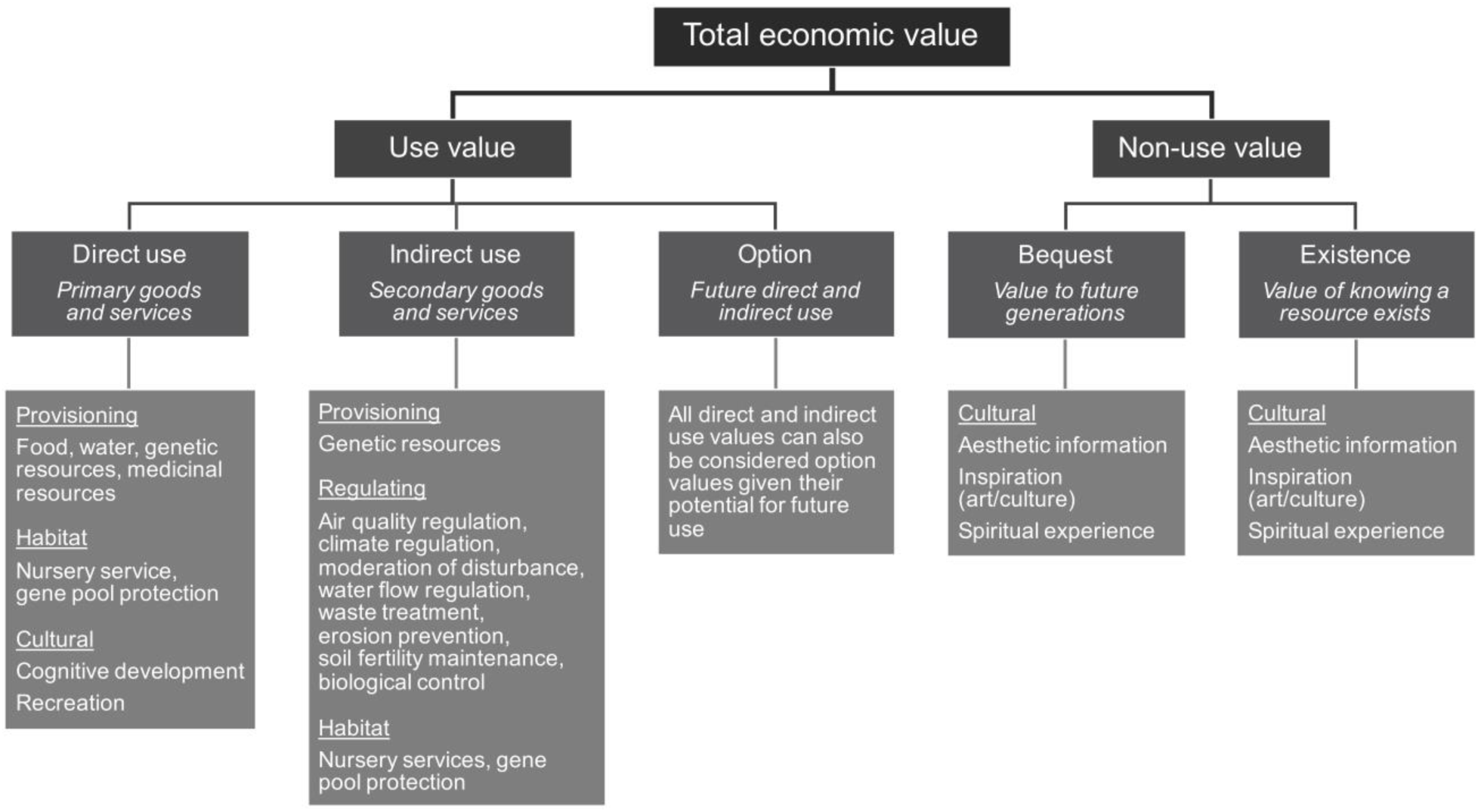
Framework modeled on TEEB (2010) and used in this paper to consider ecosystem services of the Arctic National Wildlife Refuge. Some services (e.g., ornamental resources, pollination) were omitted if there were no data to allow for valuation.

### Food

Small numbers of Alaska Native people live in and around the Arctic National Wildlife Refuge and obtain food from the local environment, a practice that has occurred for thousands of years. We therefore estimated the value of food gathered by native people from the Arctic National Wildlife Refuge that replaces commercially-purchased food. We began by calculating the population of native people in the Arctic National Wildlife Refuge region. We created a buffer that extended 80 km from the border of the Refuge and identified all of the towns inside this border. We downloaded data for 2016 city population sizes from the U.S. Census Bureau (U.S. Census Bureau 2010). We totaled the entire population of the 80-km radius and then multiplied it by the percentage of individuals in the region that identified as American Indian or Alaska Native on the U.S. National 2010 Census (U.S. Census Bureau 2010). Some wildlife travels well beyond this 80-km buffer, but we included those calculations under *Nursery Services*. It is important to note that most of the population we identified lives either within the refuge (Kaktovik) or on the boundary of the refuge (Arctic Village). These populations are highly dependent on resources from within the protected area (USFWS 2010). The most important animal resource (porcupine caribou herd) is seasonally dependent upon Refuge lands where they calve each year (Whitten et al. 1992). So, while hunting and gathering occurs both within and outside of the Refuge boundaries, the protected area is vitally important to maintaining ecosystem function throughout the broader geographic area.

To estimate the proportion of daily caloric intake provided by hunting and gathering, we utilized the list of wild food types consumed by the native Gwich’in people (e.g., caribou, waterfowl, rabbit, and berries) as a representative of local Native Alaska populations, and then identified the number of calories per 100 grams of each food source (Center for Indigenous People’s Nutrition and Environment 2008). We found the average amount (grams) consumed per day for the foods we identified (Schuster et al. 2011). We then divided the total number of calories consumed in traditional foods by 2000, an estimate of the average daily caloric intake in the U.S. (U.S. Department of Agriculture 2016).

We estimated the cost per year of food using data from a 2016 survey completed by the University of Alaska Fairbanks regarding food costs for families in Alaska (Luick 2017). We utilized the midpoint value of the cost per week per family of four and standardized this value on a per person per year basis. We then multiplied this value by the number of people in our study region who identified as Native Alaskan or American Indian in the 2010 census (U.S. Census Bureau 2010) to estimate the total cost of food for our entire population per year. We multiplied this value by the proportion of daily caloric intake provided by traditional means and divided by the number of hectares in the entire refuge. Because of the remoteness of the area, food costs on the North Slope of Alaska are likely higher than the state average, which would make our estimate conservative. However, our method of food value calculation does not take into account costs of obtaining food through hunting and gathering, including labor, materials, and equipment, costs that are difficult to calculate. More complex valuations would account for actual costs of imported food as well as costs of acquiring food locally.

We should also note that in ES valuations, estimates for climate regulation often include some measure of food production, since alteration of carbon storage capacity would presumably have an impact on that food production value (see *Climate Regulation*). Thus, using the methods of O’Garra (2017), we assumed that climate change caused by release of stored carbon would reduce food production value by 50%. Since it was adjusted by one-half, this approach prevented double counting the current food production value,

### Water

To estimate the value of water in Arctic National Wildlife Refuge, we first found the average household water costs in the state of Alaska. The Alaska Department of Environmental Conservation Division of Water reported that rural Alaskans often spend more than 5% of their monthly household income on water systems (ADEC 2016). To estimate the number of households, we multiplied the labor force participation rate for 2016 (Tiberi 2017) by the total estimated population in the two communities adjacent to the Arctic National Wildlife Refuge that obtain water directly from the ecosystems within or dependent on the Refuge (Stankiewicz 2005; ADEC 2016; Fig. 1). We multiplied annual household income by 5% to find the total dollar value of water utilized from the Refuge. This number was divided by the total hectares of the Refuge to find the value per hectare. It should be noted that by selecting the value of 5%, our estimate is conservative (ADEC 2016), but our analysis also does not account for labor and materials.

### Raw Materials

The 1980 Alaska National Interest Lands Conservation Act called for continued subsistence activities by local residents within the Arctic National Wildlife Refuge. Raw materials are presumably collected in the Refuge by members of the local indigenous communities, but data on exact quantity of materials collected are not currently available.

### Genetic Resources

To calculate genetic diversity, we estimated the number of species in the Refuge. We found species counts for plant and vertebrate species (USFWS 2011, 2017a, 2017b). Due to the high diversity of arthropods and lack of knowledge on arthropod diversity, we used a well-known order, Lepidoptera, to determine the size of all arthropod diversity. We identified the number of species in Lepidoptera, and then extrapolated the total arthropod diversity using a proportional representative comparison (Zhang 2011). Other biodiverse taxa, such as Nematoda and Bacteria, were excluded due to a lack of information on species richness. Therefore, our total estimate is conservative. To identify the potential economic value, we used the midpoint value of bio-prospecting determined by Nunes and van der Bergh (2001), where they calculated a range of monetary values for species. We then multiplied this number by the estimated number of species and divided by the number of hectares.

This method is limited by both the high uncertainty in species diversity (especially small organisms, such as prokaryotes) and value of those existing species. Individual species can be extremely valuable to humanity directly (high market value, e.g., thermophilic bacteria in Yellowstone NP; Hunter-Cervera 1998) or indirectly through keystone ecological functions (e.g., sea otters in the eastern Pacific; Estes and Palmisano 1974). It is unknown if any individual species of the Refuge fit either of those categories.

### Medicinal Resources

Medicinal resources were calculated as part of the total economic value in bio-prospecting based on Nunes and van der Burgh (2001). We assumed medicinal values were incorporated into the calculated value for *Genetic Resources*.

### Ornamental Resources

To calculate a value for ornamental resources, we searched for local products that were collected and sold for ornamental uses. Regulations within the Refuge restrict plant collection, except for subsistence purposes, and there are no data readily available to calculate the number of ornamental plants collected in the area. We also contacted Alaska state government officials to ask about studies on ornamental business performed by their commerce department, and we were informed that there have been no studies done on the subject. Therefore, while we suspect that there are some biological resources harvested for ornamental reasons, we have no data to estimate the value.

### Air Quality Regulation

It is well known that terrestrial plant cover removes air pollutants, which presumably has a causative effect on lowering health care costs (Brunekreef and Holgate 2002). The Arctic region, although relatively low in pollution sources, does have air pollution impacts from both local and remote locations (Law and Stohl 2007). Nowak et al. (2014) estimated the county level effect of tree cover on health care costs for the conterminous U.S. To estimate the value of vegetation in air quality regulation for the Refuge, we selected the midpoint value of the lower range of vegetational cover estimated by Nowak et al. (2014) and multiplied this number by the number of hectares in the Refuge. We used this midpoint value from the lower range because the Refuge is virtually treeless (except in the far southern areas), the value represents areas of the U.S. that are similar in vegetation (i.e., treeless areas of the Great Plains), and plant removal of pollutants are, therefore, likely in the lower range.

### Climate Regulation

Climate regulation by CO_2_ and CH_4_, unlike many other local regulation services, has effects on a global scale through effects on temperature, short- and long-term weather patterns, and myriad geochemical processes. Therefore, both the stored carbon and the annually sequestered carbon have an impact on global climate and are, therefore, ecosystem services that the lands of the Arctic National Wildlife Refuge provide to the globe. Using geospatial data from the Alaska Land Carbon Assessment (Genet et al. 2017), we assessed three carbon metrics for the following categories: stored soil carbon, live carbon biomass, and carbon sequestration. This approach allowed us to determine the amount of carbon stored and the avoidance cost if we were to remove the Refuge’s carbon storage capacity. For each carbon category, the number of pixels for each class within the Refuge boundary was calculated in ArcMap using the Zonal Statistics tool and these numbers were multiplied by the corresponding class value. The total values for each class were then summed to determine the total amount of carbon in the Refuge and then converted to a per hectare basis. Since loss of living plants and stored soil carbon would be a one-time event, these two values were prorated over 25 years. Release of stored carbon from permafrost is expected to consist both CO_2_ and CH_4_, although the ratios are uncertain. We used a value of 2% methane and the balance CO_2_, which we consider a conservative estimate of what proportion of the carbon would be released in this form (Schuur et al. 2015). Methane was considered to have 21 times the global climate change impact (in terms of heat trapping ability) compared to CO_2_ (IPCC 2007). This total value of carbon was based on the estimated social cost of climate change associated with increased atmospheric carbon (Tol 2008). Our valuation therefore, calculates the total global social costs if all the carbon currently stored and sequestered in the Refuge was instead present in the atmosphere.

### Moderation of Disturbance

Most of the human population that could be affected by flood waters from the Arctic National Wildlife Refuge live in the Yukon River Basin downstream from the Refuge. To calculate a value for the water control provided by the Refuge, we began by overlaying the GIS layer for the Yukon River basin over a map of the Arctic National Wildlife Refuge. This map allowed us to calculate the proportion of Yukon River that overlaps the Refuge. We then determined the value of insured properties along the Yukon River and its tributaries (FEMA 2016). We included the tributaries because the major source of flooding on the Yukon is ice jams that back up the river through its tributaries and cause it to flood (Brabets et al. 2000). Based on recent flood frequencies, we assumed a current annual flood risk of 4% (Livingston et al. 2009) and that each flood caused a 50% loss of property. From the total amount of insured property value, we took the percentage of the river basin that falls in the Refuge and multiplied it by the total insurance liability to find the value of water control. We assumed that if the Refuge did not exist as its current water control landscape, there would be a 10% increase in flood frequency and a 25% increase in intensity (Konrad 2003; Taylor et al. 2017). We divided this total estimated increase in monetary flood damage by the number of hectares in the Refuge to give us the value per hectare per year. It should be noted that there are numerous uninsured properties in the watershed (e.g., fishing and hunting cabins), so our estimates are likely conservative.

### Water Flow Regulation

Flood damage and ground water recharge were included in *Moderation of Disturbance* and *Water* categories, respectively. We did not identify any other water flow regulation services outside of these two groups.

### Waste Treatment

Wastewater treatment costs were included in calculations in *Water*.

### Erosion Prevention

We considered erosion prevention as part of *Soil Fertility Maintenance* and *Moderation of Disturbance* and, to avoid double counting, assumed the ES value for erosion prevention is included within those two calculations.

### Soil Fertility Maintenance

Soil fertility contributes to provisioning service values, so to avoid double counting and to focus on end products (Pascual et al. 2010), we did not calculate monetary values for this service. We did, however, examine three soil nutrients (nitrogen, phosphorous, potassium) that have environmental and economic value so that we could estimate how much of these materials is stored in this Arctic system. The storage of these material likely has economic benefits beyond provisioning services, but for which we had no practical way to value. For instance, the sequestered nitrogen and phosphorus reduces human-induced overabundance of these compounds in the environment.

Nitrogen (N) was estimated by considering two metrics: annual nitrogen deposition and stored soil nitrogen. For deposition, we used a dataset retrieved from EarthDATA with global terrestrial coverage indicating the amount of atmospheric nitrogen deposition in mg/m^2^/yr (Dentener 2006). The number of pixels for each category within the Refuge boundary was calculated in ArcMap using the Zonal Statistics tool and these numbers were multiplied by the corresponding class value for nitrogen deposition. The total values for each class were then summed to determine the total amount of nitrogen deposition in the Refuge and then converted to a per hectare basis. Stored nitrogen values were obtained from a spatial data set produced by Shangguan et al. (2014). The number of pixels for each category was calculated as above and summed to determine the total amount of nitrogen stored in the Refuge.

Soil phosphorus (P) amounts were determined by estimates of average soil concentrations from the world’s biomes (Xu et al. 2013). Similarly, for potassium (K), we obtained the average soil concentrations of this nutrient in soils found in the tundra biome (Sardans and Peñuelas 2015).

### Pollination

There are no crops in the region to be pollinated. We assumed the value of gathered foods that require pollination is included in *Food*.

### Biological Control

We assumed the ES values for natural biological control services provided by the ecosystem were included in *Food* production.

### Nursery Services

To calculate our value for nursery services we identified the number of waterfowl harvested in both Canada and the United States. Using distribution maps from Bird Life International’s database (BirdLife International 2018), we identified the hunted waterfowl whose breeding ranges overlapped with the Arctic National Wildlife Refuge. We then calculated the overlap of these breeding ranges with the refuge to calculate the percentage of the breeding ranges that occur in the Refuge. We applied these proportions to the number of harvested birds of each species in U.S. and Canada to identify the number of birds harvested that were likely produced (i.e., fledged) from the refuge. We identified the average amount spent per bird by using a national survey (Raftovich et al. 2017). To identify the total expenditure, we multiplied the average annual expenditure by the number of hunters. We divided this number by the proportion of days spent hunting migratory birds then divided this value by the number of waterfowl harvested in 2016-17 to get the average expenditure per bird harvested. This number was then multiplied by the number of harvested birds that were produced in the Refuge and harvested in the U.S. and Canada. We divided this by the number of hectares to develop the price per hectare. This value does not include subsistence waterfowl harvesting, which was included in *Food*.

Other species were not included in nursery services. Small numbers of fish (salmon) are produced in the Refuge and migrate elsewhere, but these data were unavailable. Caribou, which do calve in the Refuge, were included in our *Food* calculation. Polar bears, which have important, but difficult to calculate ecological and cultural values, den in the coastal plain of the Refuge, but we attempted no valuation for this species.

### Genepool Protection

We assumed value of genepool protection was calculated within sections for *Genetic Resources* and *Medicinal Resources*.

### Total Non-use Value

To calculate the value of total non-use value, we created a nationwide survey utilizing contingent valuation methods (See Appendix). This survey was created using Google Forms and distributed through Amazon’s Mechanical Turk service to 315 participants. Survey respondents were required to be U.S. citizens and above the age of 18 years old.

Each participant was initially asked if they were familiar with the Arctic National Wildlife Refuge. Those not familiar were re-directed to an unbiased educational section outlining the location, size, biodiversity, and oil/gas resources of the area. Those that were familiar with the Refuge were then asked if they supported oil and gas drilling in Arctic National Wildlife Refuge. Then participants were asked how much of their annual income they would be willing to forgo to ensure that the Refuge stayed closed to oil and gas drilling operations and for it to remain in its natural state. They were then directed to the same supplemental education material that those unfamiliar with the Refuge were shown. After the passage, we asked if the participants supported oil and gas drilling in the Refuge and how much of their annual income, if any, they would be willing to forgo to ensure that the Refuge remained closed to oil and gas drilling. We did not attempt to differentiate why survey participants valued the Refuge in its natural state, but assumed that those reasons encompassed all non-use values (e.g., aesthetic, spiritual, art, culture, among others, both current and future).

The value utilized in our final calculations was the initial value of those familiar with the Refuge. The average willingness to forgo income was applied to the percentage of those familiar with the Refuge. The percentage of participants familiar with the Refuge was applied to the entire U.S. population of voting age, while we assumed that the percentage not familiar with the Refuge would have a $0 willingness to forgo income. We then extrapolated the average U.S. citizen’s willingness to forgo income to the entire U.S. population. To estimate what the total value could be under a higher level of public awareness, we also reported the value after the short educational information had been received (using the same methods as above).

### Aesthetic Information

We assumed that the aesthetic value of the Refuge was incorporated into the survey completed in the section for *Total Non-use Value*.

### Inspiration for culture and art

We assumed inspiration for culture and art was incorporated into values obtained from the survey in the section for *Total Non-use Value*. Local Native Alaskan people (e.g., Gwich’in) place a high cultural value on the landscape of the Refuge and surrounding lands (Parlee and Berkes 2005). Since the culture of these populations is so dependent on the local environment and would radically change without the resources of the Refuge, it is also difficult to monetize this value.

### Spiritual Experience

We assumed spiritual values were included in the survey for *Total Non-use Value.* As noted in previous section, many local Native Alaskan people are likely to view the landscape as having extraordinary spiritual value, but we find this value incalculable.

### Cognitive Development

To develop a value for cognitive development, we utilized the annual park budget as well as funds spent on research within the Refuge. To calculate the value spent to protect the gene pool in Arctic National Wildlife Refuge, we used the total value of the 2016 fiscal year budget for the Refuge, which was provided by a refuge staff member (J. Reed, personal communication). We then divided the total budget value by the number of hectares in the refuge to identify the value per hectare. For research information, we contacted the Refuge and requested information on all research projects being performed in the Refuge. We identified ten research projects taking place in the Refuge and contacted each researcher attached to each of these projects. We received responses from eight out of the ten researchers and recorded their responses for their total budget. To develop an annual budget, we divided the total budget by the number of the years they will be working and added all of the annual budgets together. With this sum of the money spent annually on research in the Refuge, we divided by the total number of hectares in the Refuge to calculate the value per hectare per year.

### Recreation

We assumed recreation was both a consumptive (e.g., hunting) and a non-consumptive (e.g., hiking) direct use value (Pascual et al. 2010). We utilized recreation expenditures as a proxy for recreational value. To calculate this value for recreation each year, we first identified the number of visitors to the park in the year 2016 by directly contacting refuge staff. We utilized a public use report released by the U.S. Fish and Wildlife Service that gave us the percentages of visitors who came for hunting and those who came for non-hunting recreational activities (USFWS 2010). We surveyed 11 different outfitters and determined the average cost of 29 different recreational trips offered. We repeated this method for every hunting guide who had legal access into the Refuge. We surveyed guide services with public prices and averaged the price of 21 guided hunts and 17 trophy fees. Based off the public use survey, every hunter takes a trophy with their hunt. We then multiplied the number of visitors by the percentage of recreation usage and the proportion who came for hunting. We then multiplied the individual population numbers times the average cost of the respective trips. After summing these numbers, we then divided the total money spent on the ground by the number of hectares in that park to get our specific value per hectare. In addition, we used a visitor study (Christensen et al. 2017) to determine the home location of visitors and then found the average cost of a flight from the major airport in the area of their home state to the Fairbanks airport. To calculate the average cost of an international flight, we determined the average cost of a flight to Alaska from the seven largest airports outside the U.S. Flights from major cities in Alaska to recreational locations are included in guide packages (described above). All of these average travel costs were then added together and divided by the number of hectares in the Refuge to calculate the average travel cost per hectare.

### Detailed examination of 1002 Area

Since there is controversy over oil drilling in the 1002 Area, we performed an analysis of this specific region for select variables that differ geographically within the Refuge and for which we had detailed spatial data that allowed us to analyze this variation. Both carbon and nitrogen values have GIS layers readily available with enough resolution to discern variation within the Refuge, so we calculated the ES values for *Climate Regulation* and amounts of nitrogen for *Soil Fertility* using the same methods described above. Since we originally used broad estimates for phosphorus (P) and potassium (K) and there are no spatial data available, we did not adjust them for the 1002 Area, so only the variation in nitrogen values affected the soil fertility calculations. The climate regulation values were added to the broad estimates for the other ES categories to calculate a total for the 1002 Area while the soil fertility amounts are provided for informational purposes.

## Results

### Overview

Our estimates place an annual value of about 13 billion USD for the Arctic National Wildlife Refuge, for a per hectare value of 1,708.94 USD (Table 1). The majority of these ES values originate from two categories: climate regulation and non-use values, which together account for 99% of ecosystem services. These two are explained in detail below. In addition, we have also included a detailed description of the amounts for soil fertility maintenance in the Refuge because these stored amounts contribute to other important ecosystem services (e.g., provisioning and aesthetic information).

**Table 1.**
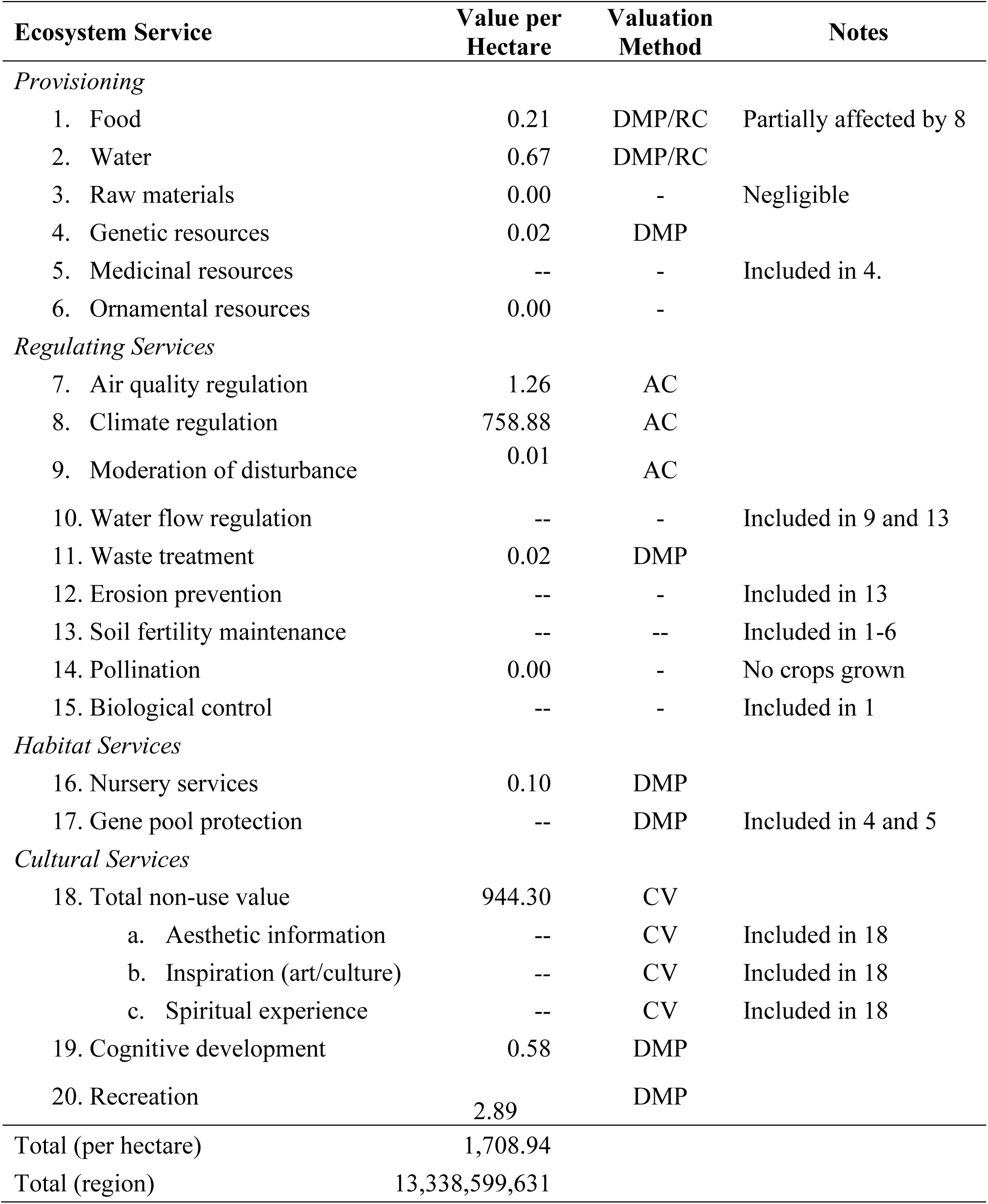
Annual ecosystem services values estimated for the Arctic Refuge in 2016 USD. DMP = direct market price, RC = replacement cost, AC = avoidance cost, CV = contingent valuation.

#### Climate regulation

Over 90% of the ES value for carbon in the Arctic National Wildlife Refuge can be attributed to carbon stored in the permafrost of the soil, mostly as carbon dioxide (CO_2_) with smaller amounts of methane (CH_4_, Table 2). Methane is a relatively small proportion of the stored carbon (about 2%), but because of the increased global climate change impacts of methane (about 21 times greater than CO_2_), it makes up about one-third of the stored carbon ES value. As a low productivity environment in a higher latitude, smaller amounts of carbon are stored as live biomass (Table 2). This region continues to act as a carbon sink, although the annual amount of carbon per hectare stored is small. There is also much variation in soil sequestration, with some areas absorbing larger amount of carbon each year and other areas acting as carbon sources (Fig. 3).

**Fig. 3.**
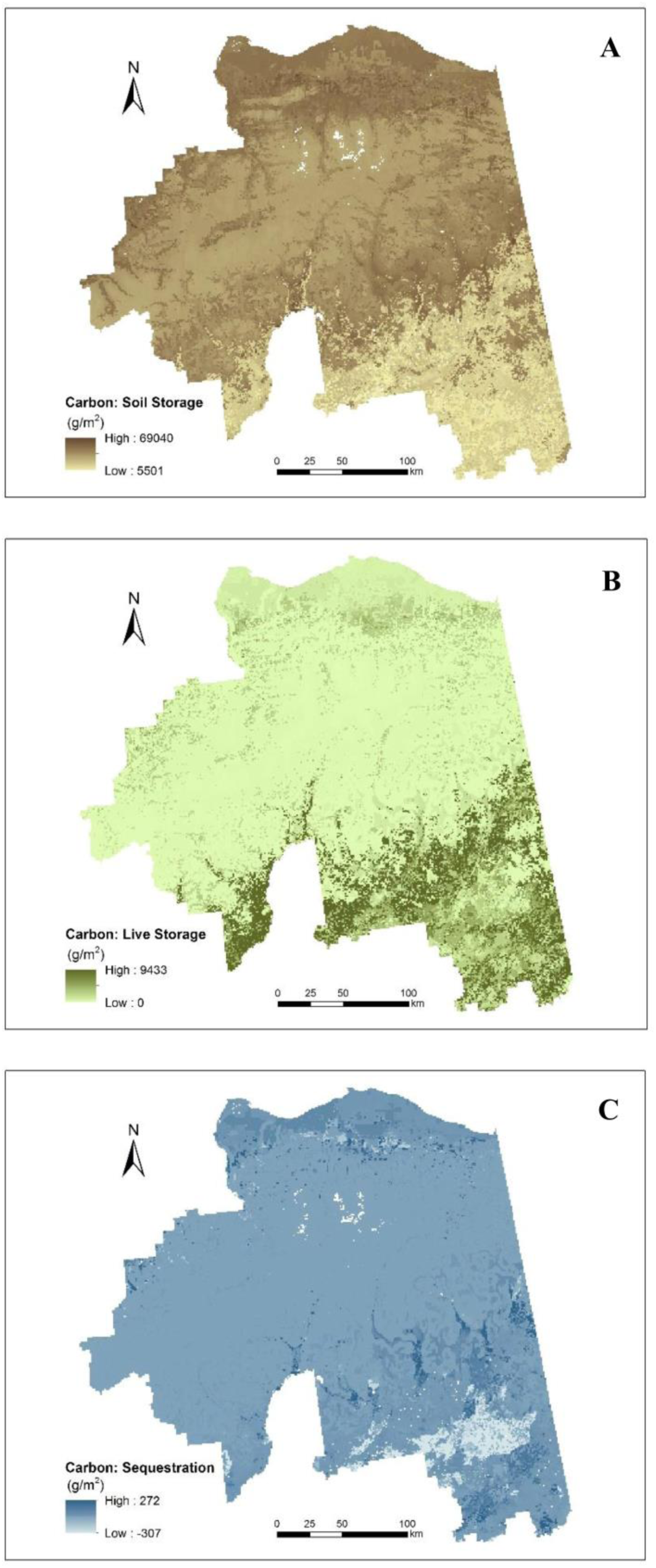
Spatial distribution of variables used to calculate the ecosystem services value for climate regulation in the Arctic Refuge. Variables include the amount of land carbon (g/m^2^) assessed over the time period from 2000-2009 that is a) stored in the soil, b) stored in live biomass, and c) sequestered annually. Data source: USGS 2017 Alaska Land Carbon Data Assessment.

**Table 2.** Total amounts and ecosystem services (ES) values of carbon and amounts of select nutrients stored in the Arctic Refuge. Note that ES values for sources stored in soil and live biomass are prorated over 25 years. Values given in 2016 USD. Ecosystem services of the Arctic NWR

| Entire Refuge |  |  |  |  |  |
| --- | --- | --- | --- | --- | --- |
| Source | Amount per Hectare |  |  |  |  |
|  | Carbon<br>(tonnes in CO<sub>2</sub> form) | Carbon<br>(tonnes in CH<sub>4</sub> form) | Nitrogen<br>(tonnes) | Phosphorus<br>(tonnes) | Potassium<br>(tonnes) |
| Live biomass | 32.26 | - | - | - | - |
| Stored in soil | 337.48 | 6.89 | 18.40 | 0.24 | 0.46 |
| Annual sequestration | 0.03 | - | 0.0003 | - | - |
| Annual Ecosystem Service Value per Hectare |  |  |  |  |  |
|  | Carbon (CO<sub>2</sub> form) | Carbon (CH<sub>4</sub> form) | Nitrogen | Phosphorus | Potassium |
| Live biomass | 47.53 | - | - | - | - |
| Stored in soil | 497.18 | 213.08 | - | - | - |
| Annual sequestration | 1.10 | - | - | - | - |
| 1002 Area |  |  |  |  |  |
| Source | Amount per Hectare |  |  |  |  |
|  | Carbon<br>(tonnes in CO<sub>2</sub> form) | Carbon<br>(tonnes in CH<sub>4</sub> form) | Nitrogen<br>(tonnes) | Phosphorus<br>(tonnes) | Potassium<br>(tonnes) |
| Live biomass | 6.29 | - | - | - | - |
| Stored in soil | 350.91 | 7.16 | 19.40 | 0.24 | 0.46 |
| Annual sequestration | 0.21 | - | 0.0027 | - | - |
| Annual Ecosystem Service Value per Hectare |  |  |  |  |  |
|  | Carbon (CO<sub>2</sub> form) | Carbon (CH<sub>4</sub> form) | Nitrogen | Phosphorus | Potassium |
| Live biomass | 9.26 | - | - | - | - |
| Stored in soil | 516.97 | 221.56 | - | - | - |
| Annual sequestration | 7.73 | - | - | - | - |

#### Non-use Valuation

Willingness to pay values were utilized to estimate non-use cultural values (aesthetic, spiritual, inspiration for art and culture) and preference for protection from a survey of U.S. citizens (Table 3). Surprisingly, only 47% of the survey participants were aware of the Arctic National Wildlife Refuge, and so a relatively small proportion of people were able to give a contingent valuation. However, a large majority who were aware of the Refuge did not support oil exploration and drilling and were willing to forego a considerable amount of income, which produced a large monetary value. When given a short, neutral, educational statement about the Refuge, a slightly smaller, but still considerable majority of the survey participants (which now included all) supported maintaining the Refuge in its natural state. Using this value would increase the contingent valuation by more than 78% and increase total ES values for the Arctic National Wildlife Refuge by about one-third. However, we retained the conservative value (before educational information) for our comprehensive ES estimate (Table 1) since we believe it more accurately represents the current state of the contingent valuation for U.S. population.

**Table 3.** Results of the contingent valuation survey for the Arctic National Wildlife Refuge. Survey details described in Appendix I. N = 315 participants.

|  |  | Familiar with Arctic National Wildlife Refuge (47%) |  |  | Not familiar with Arctic National Wildlife Refuge (53%) |  |  | Weighted Mean | Value per hectare | Total value |
| --- | --- | --- | --- | --- | --- | --- | --- | --- | --- | --- |
|  |  | Support drilling | Do not support drilling | No opinion | Support drilling | Do not support drilling | No opinion |  |  |  |
| Before education | Percentage of respondents | 17.69% | 73.47% | 8.84% | - | - | - | - | - | - |
| | Willingness to pay per person | \$0 | \$90.01 | \$42.69 | - | - | - | \$84.97 | \$944.30 | \$7.37 bil. |
| After education | Percentage of respondents | 23.13% | 71.43% | 5.44% | 19.64% | 68.45% | 11.90% | - | - | - |
| | Willingness to pay per person | \$0 | \$98.39 | \$51.88 | \$0 | \$65.97 | \$14.75 | \$75.20 | \$1,687.19 | \$13.2 bil. |

### Soil Fertility Maintenance

The three major limiting nutrients of the soil estimated (N, K, P) are abundant in this system (Table 2). It should be noted that these nutrients help promote ecosystem function in the region, although provisioning services are relatively small due to the small human population benefitting from these services. Due to the remoteness and harsh environment, the nutrients are not readily accessible to human activity or benefit, but the Refuge may create economic benefits by preventing their release into the environment.

## 1002 Area

Since we had no way to determine the variation in most ES across the Refuge, we assumed values other than carbon were a constant value per hectare regardless of location. When considering the variation in carbon, comprehensive ES values for the 1002 Area were estimated at 1705.79 USD. This value is slightly lower compared to the corresponding value for the Refuge as a whole. While the amount of carbon stored in the soil and amount sequestered was higher in this region, the amount of live biomass was considerably lower (Table 2).

## Discussion

The overall estimated value of 1,708.94 USD/ha/year is lower than most other terrestrial biomes (Costanza et al. 2014), although higher than the desert biome (Taylor et al. 2017). Two ES categories (climate regulation and total non-use value) made up over 99% of the total value. This result is not entirely surprising since Arctic tundra lacks some of the resources that tend to increase other terrestrial biomes values, such as food production and raw materials (e.g., wood). Additionally, human population density in the region is extremely small, so that local ES values are relatively small, albeit not for the local individuals relying upon these ES. However, the Arctic tundra provides important global values, in particular the two highest values: total non-use value and climate.

While the local ES estimates are relatively small in direct monetary value, it is important to emphasize that the process of ES valuation is noted for its difficulties in accurately estimating cultural values (Chan et al. 2012). Some groups of Native Alaskans have been strongly opposed to oil and gas drilling in the region (e.g., Gwich’in, Krech 2005). Many of their concerns focus on the negative impacts of development in a landscape that is viewed as sacred. The unquantifiable aspects the Refuge maintains for cultural and spiritual well-being should, therefore, not be disregarded.

Kotchen and Burger (2007) estimated the mean economic benefit of 1,141 (2005 USD) per person of voting age over the 30-year period of expected oil production (about 38 USD per person per year). This economic impact of oil development corresponds to a 1,378.62 (2016 USD) per hectare per year for the refuge. Our willingness-to-pay results from our survey indicated that, based on current awareness of the Arctic National Wildlife Refuge issues, the U.S. population is willing to pay less (68%) in lost income than the value of the oil (Table 3).

However, this value is based on over one-half of the population having no knowledge of the refuge, a surprisingly high number considering the amount of media attention the region has garnered over the last two decades (Bengston et al. 2010). When the survey cohort was provided with neutral educational information and we were then able to include all participants in the value estimation, the total willingness-to-pay value rose to 1,687.19 (2016 USD) per hectare per year, which is 22% higher than the estimated oil value. Therefore, we argue that the U.S. public is currently undereducated about the Arctic National Wildlife Refuge and that increased information about the oil drilling controversy is likely to change the number of people concerned about fossil fuel development versus continued protection. While people who already have a firmly held position on drilling in the Refuge are unlikely to change their view because of new information (Teel et al. 2006), our results suggests that since a large portion of the population has no knowledge of the controversy, an education program could be impactful. Therefore, a public education program aimed at increasing the wilderness and conservation perceptions of the Refuge could change political pressure to restrict future energy development.

Our results indicate that the ES value of the 1002 Area is not higher compared the Refuge as a whole. The coastal plain region of the Refuge is probably more valuable for certain wildlife (e.g., waterfowl, caribou, polar bear), but these provide relatively small amounts of ecosystem services compared to other categories. The one caveat is that, in our survey, we did not distinguish the non-use value of the coastal plain, which, considering the charismatic wildlife dependent in that region, might be higher among U.S. residents.

The climate regulation value is important for this region, and Arctic tundra in general, because of the large amount of carbon stored as CO_2_ and CH_4_ in the permafrost. Arctic tundra has one of the largest amounts of soil sequestered carbon of any biome on Earth. However, the warming climate is putting this ES at risk. As tundra warms, permafrost is melting over large areas of the Arctic, releasing these greenhouse gases in increasing amounts (Schuur et al. 2009). Models predict that the tundra will be a net emitter of carbon by the mid-21^st^ Century, although the exact date is uncertain (Koven et al. 2011; Schaefer et al. 2011). Regardless of whether energy is extracted from this region, the carbon sequestration value will be negative for the entire Refuge, although the stored carbon will take many decades to be released into the atmosphere.

Therefore, it is important to note that the current value of climate regulation this environment possesses is likely dependent on the broader human culture to solve the current greenhouse gas emissions crisis.

Since the passage of U.S. Bill H.R.1 2018, the Arctic National Wildlife Refuge has been undergoing an Environmental Impact Statement process regarding the potential effect of oil drilling on the 1002 Area of the Refuge (BLM 2018, USDI and BLM 2019). Leases will presumably be offered in the near future, although legal actions are expected to delay drilling for some time (Grandoni and Eilperin 2020). If drilling does indeed proceed, impacts on the environment are likely to be pervasive. Fossil fuel development has large impacts on land-use, ecosystem services, landscape fragmentation, and wildlife (Jones and Pejchar 2013; Thompson et al. 2015; Moran et al. 2015; 2017; Allred et al. 2015; Trainor et al. 2016). Any environmental impact analysis needs to include ecosystem services loss as one the externalities associated with oil drilling. The loss of non-use valuation could be particularly pronounced if the wilderness characteristic of the coastal plain of the Refuge is lost. Unlike other important protected areas that have tourist infrastructure and are commonly visited by the humans (e.g., Yellowstone National Park), the Arctic National Wildlife Refuge is unlikely to be a major destination for tourists, mostly because of costs and limited accessibility. However, our survey results indicate that the public strongly values this landscape and would be willing to forego income greater (if educated) than the value of any resource extraction, indicating support for oil drilling is limited and that the damage could reduce its long-term ES value. This result shows the value people place on remote and undeveloped parts of the world, which includes much of the greater Arctic region.

## Appendix. Structure of the contingent valuation survey provided to a selection of U.S. residents.

Arctic National Wildlife Refuge National Survey (5 minutes) Please only take this survey if you are 18 years or older.

1. Are you 18 years or older?
  a. Yes [directed to part 2]
  b. No [survey ends]
2. Are you familiar with the Arctic National Wildlife Refuge (Arctic National Wildlife Refuge)?
  a. Yes [directed to part 3]
  b. No [directed to part 5 Arctic National Wildlife Refuge description]
3. Do you support oil and gas drilling in the Arctic National Wildlife Refuge (Arctic National Wildlife Refuge)?
  a. Yes [directed to part 4]
  b. No [directed to part 4]
  c. Don’t Know/No Opinion [directed to part 4]
4. Considering your annual income, how much money would you be willing to give up per year in order to continue a ban on oil and gas drilling in the Arctic National Wildlife Refuge (Arctic National Wildlife Refuge)? If you select “other”, please enter an amount in US Dollars ($).
  a. $50 [directed to part 5]
  b. $100 [directed to part 5]
  c. $200 [directed to part 5]
  d. $500 [directed to part 5]
  e. Other: [directed to part 5]
5. Read Description of Arctic National Wildlife Refuge
  The Arctic National Wildlife Refuge is one of the largest undeveloped areas in North America. It has the most species of any similar size area in the Arctic, including one of the largest caribou herds in the world. Currently, it is fully protected from development and human activities other than recreation. The area is also believed to contain a large oil and gas deposit. The Tax Cuts and Jobs Act passed by Congress in 2017 opened about 8% of the Arctic National Wildlife Refuge to oil and gas drilling. It is expected that this portion of the refuge will produce an estimated 7.7 billion barrels of oil over 30 years, which represents about 5% of the expected US total production. Some Americans believe that the area should remain undeveloped as a nature preserve, while others believe the US should utilize the oil and gas deposits in the refuge.
6. Now that you have read the previous paragraph, do you support oil and gas drilling in the Arctic National Wildlife Refuge (Arctic National Wildlife Refuge)?
  a. Yes [directed to part 7]
  b. No [directed to part 7]
  c. Don’t Know/No Opinion [directed to part 7]
7. Considering your annual income, how much money would you be willing to give up in order to continue a ban on oil and gas drilling in the Arctic National Wildlife Refuge (Arctic National Wildlife Refuge)? If you select “other”, please enter an amount in US Dollars ($).
  a. $50 [survey ends]
  b. $100 [survey ends]
  c. $200 [survey ends]
  d. $500 [survey ends]
  e. Other: [survey ends]

## Acknowledgements

Thanks to the staff of the Arctic National Wildlife Refuge for their help in data collection. L. Kennedy assisted with development of the contingency valuation survey. T. Fullman, N. Whittington-Evans, and D. Krause read early drafts of this manuscript and provided insightful feedback.

